# U-Infuse: Democratization of Customizable AI for Object Detection

**DOI:** 10.1101/2020.10.02.323329

**Authors:** Andrew Shepley, Greg Falzon, Christopher Lawson, Paul Meek, Paul Kwan

**Author notes:** **Correspondence**: Andrew Shepley (alternative).

## Abstract

1. Image data is one of the primary sources of ecological data used in biodiversity conservation and management worldwide. However, classifying and interpreting large numbers of images is time and resource expensive, particularly in the context of camera trapping. Deep learning models have been used to achieve this task but are often not suited to specific applications due to their inability to generalise to new environments and inconsistent performance. Models need to be developed for specific species cohorts and environments, but the technical skills required to achieve this are a key barrier to the accessibility of this technology to ecologists. There is a strong need to democratise access to deep learning technologies by providing an easy to use software application allowing non-technical users to custom train custom object detectors.
2. U-Infuse addresses this issue by putting the power of AI into the hands of ecologists. U-Infuse provides ecologists with the ability to train customised models using publicly available images and/or their own camera trap images, without the constraints of annotating and pre-processing large numbers of images, or specific technical expertise. U-Infuse is a free and open-source software solution that supports both multiclass and single class training and inference, allowing ecologists to access state of the art AI on their own device, customised to their application without sharing IP or sensitive data.
3. U-Infuse provides ecological practitioners with the ability to (i) easily achieve camera trap object detection within a user-friendly GUI, generating a species distribution report, and other useful statistics, (ii) custom train deep learning models using publicly available and custom training data, (iii) achieve supervised auto-annotation of images for further training, with the benefit of editing annotations to ensure quality datasets.
4. Broad adoption of U-Infuse by ecological practitioners will improve camera trap image analysis and processing by allowing significantly more image data to be processed with minimal expenditure of time and resources. Ease of training and reliance on transfer learning means domain-specific models can be trained rapidly, and frequently updated without the need for computer science expertise, or data sharing, protecting intellectual property and privacy.

## I. Introduction

The use of camera trap image analysis in biodiversity management is one of the primary means by which ecological practitioners monitor wildlife (O’Connell, Nichols et al. 2011, Bengsen, Robinson et al. 2014, Meek, Fleming et al. 2014, Rovero and Zimmermann 2016, Lashley, Cove et al. 2018), obtain species distribution (Li, Bleisch et al. 2018), perform population estimates (Ahumada, Hurtado et al. 2013, Zaumyslova and Bondarchuk 2015, Fegraus and MacCarthy 2016, Rahman, Gonzalez et al. 2016) and observe animal behavioural patterns (Rowcliffe, Kays et al. 2014). However, this results in millions of images being captured, which must be processed. This is a time and resource expensive task, often manually undertaken by ecologists, which has given rise to a strong need for automation (Falzon, Meek et al. 2014). This has triggered significant interest in deep learning-based image processing solutions (Swanson, Kosmala et al. 2015, Gomez Villa, Salazar et al. 2016, Norouzzadeh, Nguyen et al. 2018, Schneider, Taylor et al. 2018, Willi, Pitman et al. 2018, Miao, Gaynor et al. 2019, Tabak, Norouzzadeh et al. 2019, Falzon, Lawson et al. 2020).

Wildlife Insights (https://wildlifeinsights.org) provides an online cloud-based service allowing practitioners to upload camera trap images to a Google Cloud-based platform which filters empty images and performs classification on 614 species. Similarly, Microsoft AI for Earth Camera Trap API (https://github.com/microsoft/CameraTraps) uses a cloud-based system to perform object detection on large quantities of camera trap images using MegaDetector. It can be used in conjunction with TimeLapse2 (http://saul.cpsc.ucalgary.ca/timelapse/) and Camelot (https://gitlab.com/camelot-project/camelot), however inference is only available for separation of empty/non empty images, and detection of limited classes. Other cloud-based inference systems include Project Zamba (https://zamba.drivendata.org/), which is a Python toolkit specific to African species and Conservation AI (https://conservationai.co.uk/), which provides access to 4 UK and Africa models upon registration. Although these services facilitate the task of camera trap image processing, users do not have the option to train models on their own images, nor can they use the services on their own device, without sharing their image data. Furthermore, models are often not sufficiently location invariant to be used with high confidence, limiting their widespread usage.

Alternatives to cloud-based services include ClassifyMe (Falzon, Lawson et al. 2020), which upon registration provides offline, on-device access to more than five YOLOv2 (Redmon and Farhadi 2016) models trained on publicly available camera trap datasets. However, these models are highly optimised for specific environments, meaning they do not generalise well to unseen environments, and users are unable to train their own models. Another alternative is Machine Learning for Wildlife Image Classification (MLWIC) (Tabak, Norouzzadeh et al. 2019), which allows the development of custom models using the R Programming Language (Team 2006). However, this requires technical knowledge and a large investment of time and resources into model development. Similarly, Camera-Trap-Classifier (Willi, Pitman et al. 2018), which is an experimental camera trap object detector, requires knowledge of Unix, limiting its adoption by non-technical ecological practitioners.

One significant issue faced by all object detection solutions is the lack of location invariance of deep learning models, and their inability to generalise to unseen environments (Yu, Jiangping et al. 2013, Beery, Van Horn et al. 2018, Miao, Gaynor et al. 2019). Due to the inherent difficulty represented by high occlusion, illumination, high object density, camouflage, movement and poor data quality usually featured in camera trap images, the development of high precision camera trap object detectors capable of generalisation to any environment is a challenge that is yet to be achieved (Gomez Villa, Salazar et al. 2016). This means they lack sufficient accuracy to be deployed in domains not included in the training data (Miao, Gaynor et al. 2019). Transfer learning has been used to improve ability to generalise (Willi, Pitman et al. 2018) however optimal performance can only be attained if the ecological practitioner has access to a model trained on their own data. Furthermore, development and deployment of such models requires specialised computer science skills (Falzon, Lawson et al. 2020). This need is addressed by U-Infuse, which provides a means by which ecological practitioners can easily train their own deep learning object detectors without requiring significant investment of time in the technical programming and specialised artificial intelligence domain knowledge required to develop custom models. U-Infuse is designed to enable ecologists to achieve location invariant object detection on camera trap images according to the methodology proposed by (Shepley, Falzon et al. 2020). This would allow practitioners to process image data in-house, at their own pace according to their needs, removing the need to allocate significant time and resources to upload, save and share data with service providers which is a drawback of existing solutions.

## II. The U-Infuse Application

U-Infuse is a free, open-source software application supported by Windows 10, Linux and macOS operating systems. The U-Infuse app is developed in the Python 3 programming language with bindings to core Python-based machine learning frameworks and image processing facilities along with a Qt5 Graphical User Interface (GUI). It has been verified on Windows 10 Home and Professional, Ubuntu 18.04-20.04, and CentOS. Copying, distribution and modification of U-Infuse source code is encouraged. Accordingly, U-Infuse is distributed under the terms of a GNU General Public License (https://www.gnu.org/licenses/gpl-3.0.en.html).

The simple-to-use GUI allows users to auto-annotate training images, custom train their own object detectors using RetinaNet (Lin, Goyal et al. 2018), and perform inference using pretrained or custom models on custom datasets. GUI Performance has been verified for large datasets, containing approximately 10,000s of images. U-Infuse is available online at GitHub (https://github.com/u-infuse/u-infuse). All U-Infuse functionalities are also provided via Python scripts and Jupyter Notebooks, which contain the U-Infuse pipeline that can be used as is, incorporated into, or adapted to any project on any platform. Whilst the GUI is appropriate for workstations processing thousands of images, the Jupyter Notebooks allow the U-Infuse functionalities to be extended to datasets of any size, on HPC systems. All upgrades, demonstrations and tutorials are available on the corresponding GitHub Wiki.

Installation of U-Infuse is straightforward and is achieved either from source code or via a downloadable binary complete with an installation wizard (Windows only). Software dependencies include RetinaNet (Lin, Goyal et al. 2018), TensorFlow (Abadi, Barham et al. 2016), OpenCV and Python 3. The installation script automatically incorporates these dependencies within the installation. Model training via the Graphical Processor Unit is facilitated for CUDA supported hardware and requires cuDNN and the CUDA development toolkit (which must be installed by the user).

### a) Functionalities

U-Infuse provides users with a complete object detection pipeline, supporting image annotation, object detector training and inference capabilities. It uses as input image datasets supporting all the most commonly used image formats including PNG, JPEG and TIFF, and optional corresponding annotation files (.xml format). All training scripts and files are contained within the U-Infuse installation, or generated by user-initiated training, inference or annotation processes. See Figure 1 for workflow diagram.

**Figure 1:**
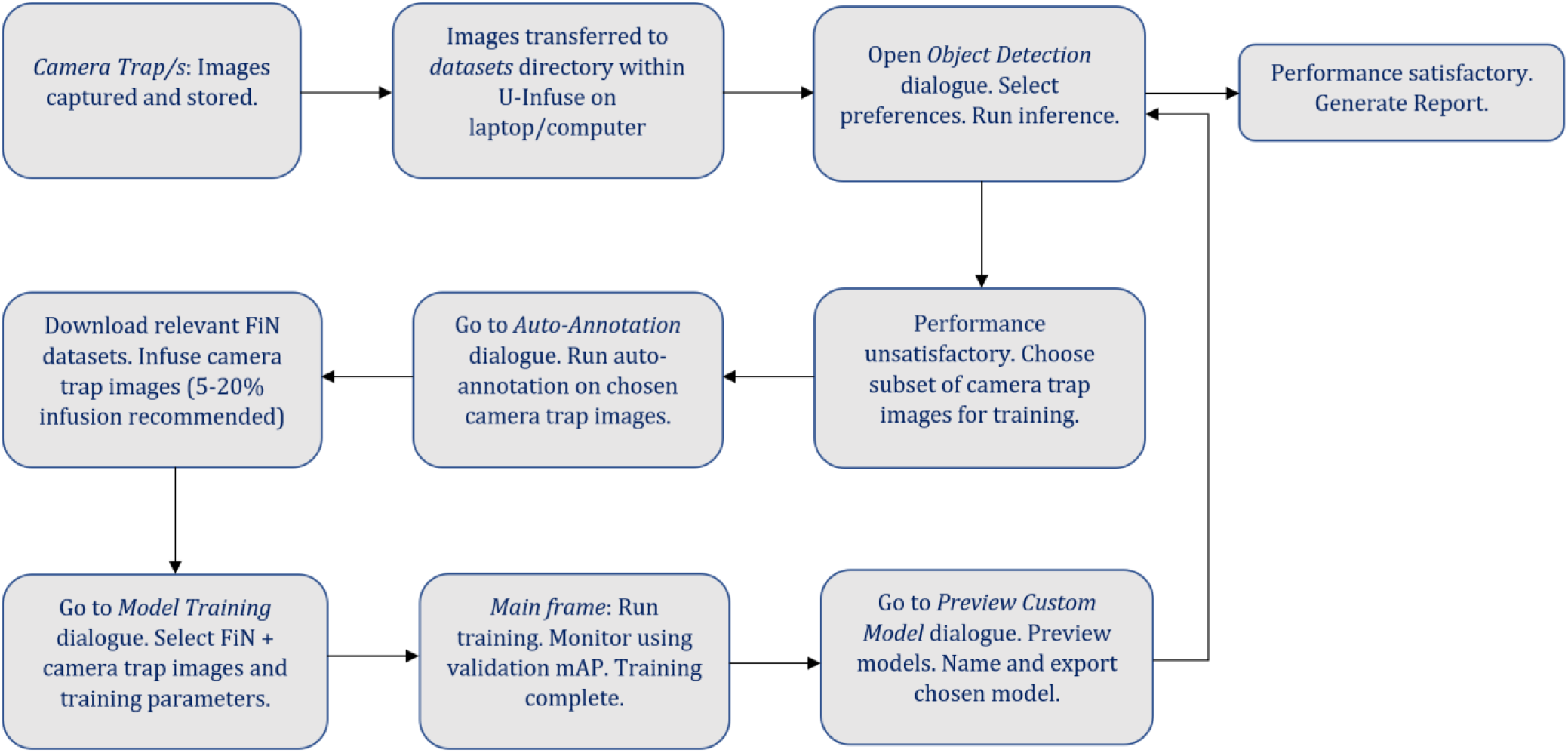
Sample workflow diagram demonstrating pipeline useability. FiN refers to FlickR and iNaturalist data as described in (Shepley, Falzon et al. 2020).

## IV. Animal Detection and Classification Using Default Models

The U-Infuse installation comes with six pretrained RetinaNet object detector models. Detailed information about these models are provided in Table 1. These models are provided to be used as pretrained models for user-controlled transfer learning, for inference and demonstration purposes.

**Table 1:** Summary of pretrained models included with U-Infuse installation. See Appendix 1 for available classes.

| Model Name | Classes | Source |
| --- | --- | --- |
| *COCO_pretrained* | 80 classes | MS COCO |
| *Australian_Multi-class* | 30 classes | FlickR |
| *pig_single_class* | Pig | FlickR |
| *striped_hyena_single_class* | Dog | FlickR |
| *rhino_single_class* | Cat | FlickR |

These models can be used for camera trap image classification via the *Object Detection* dialogue, shown in Figure 2. Users can select the model of their choice via the dropdown list (1). The chosen model will perform object detection on all the images contained in the chosen image folder (2).

**Figure 2:**
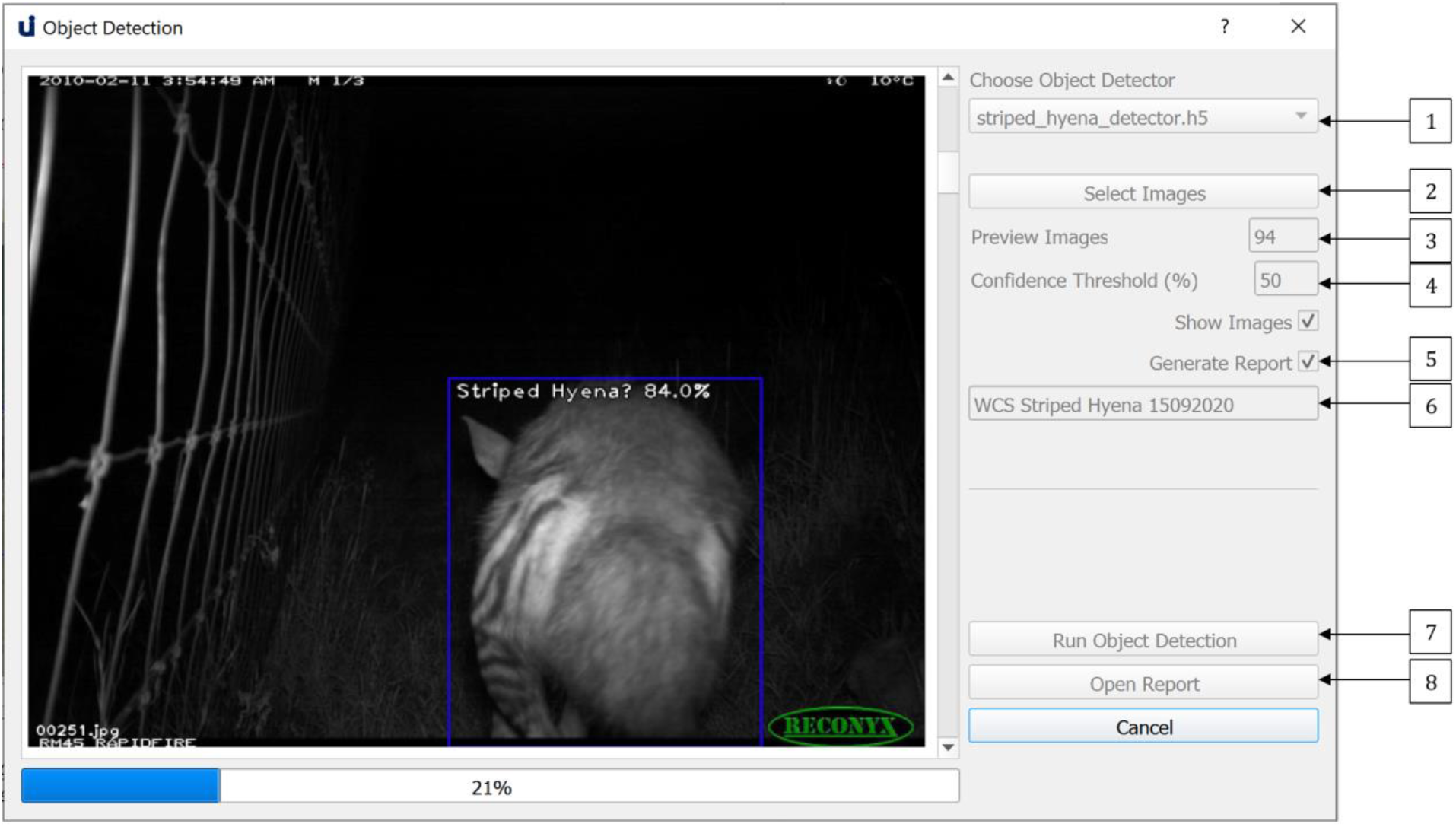
The Object Detection dialogue allows users to perform object detection on a dataset of images, optionally generating an object detection report

Users may optionally limit the number of images on which inference is conducted via the option at (3), with the default being the number of images in the chosen folder. After choosing the level of desired accuracy via the confidence threshold (4), they may elect to show images while inference is running and generate an inference report (5).

If they choose to generate an inference report, they must provide a name for their report (6). A summary and detailed report will be generated in JSON format and saved in the *reports* directory. The summary report contains information about the model used, the dataset, and the object detection output, such as species distribution and number of empty images. The detailed report contains object detection data for each image, for example, number of objects, class, confidence of detections and bounding box coordinates. Users can optionally generate a JSON file containing references to each empty image. A sample of the Summary Report is shown in Figure 3, and a sample of the Detailed Report is shown in Figure 4. Figure 3. The Summary Report can be opened via the open at (8).

**Figure 3:**
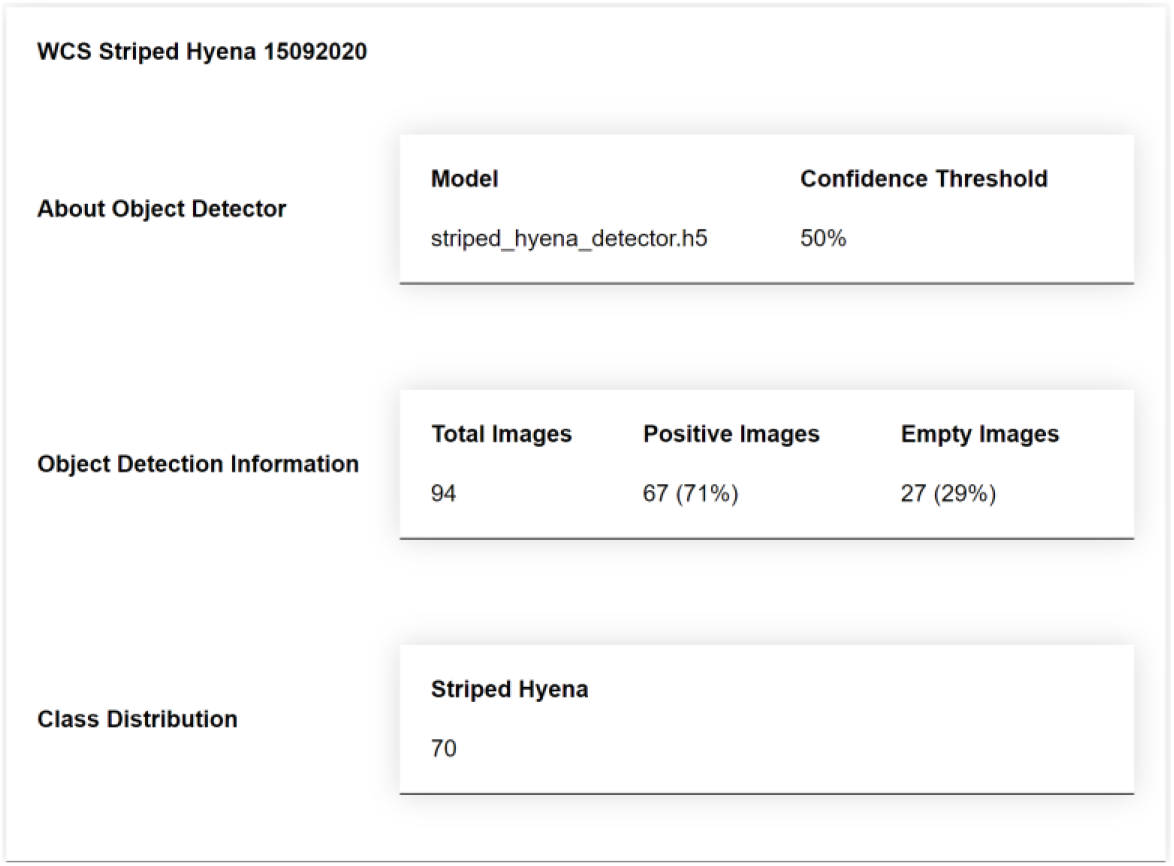
A sample Summary Report generated by running inference on a dataset of images. The report provides a summary of object detection information, including class distribution, number and percentage of positive images compared to empty images.

**Figure 4:**
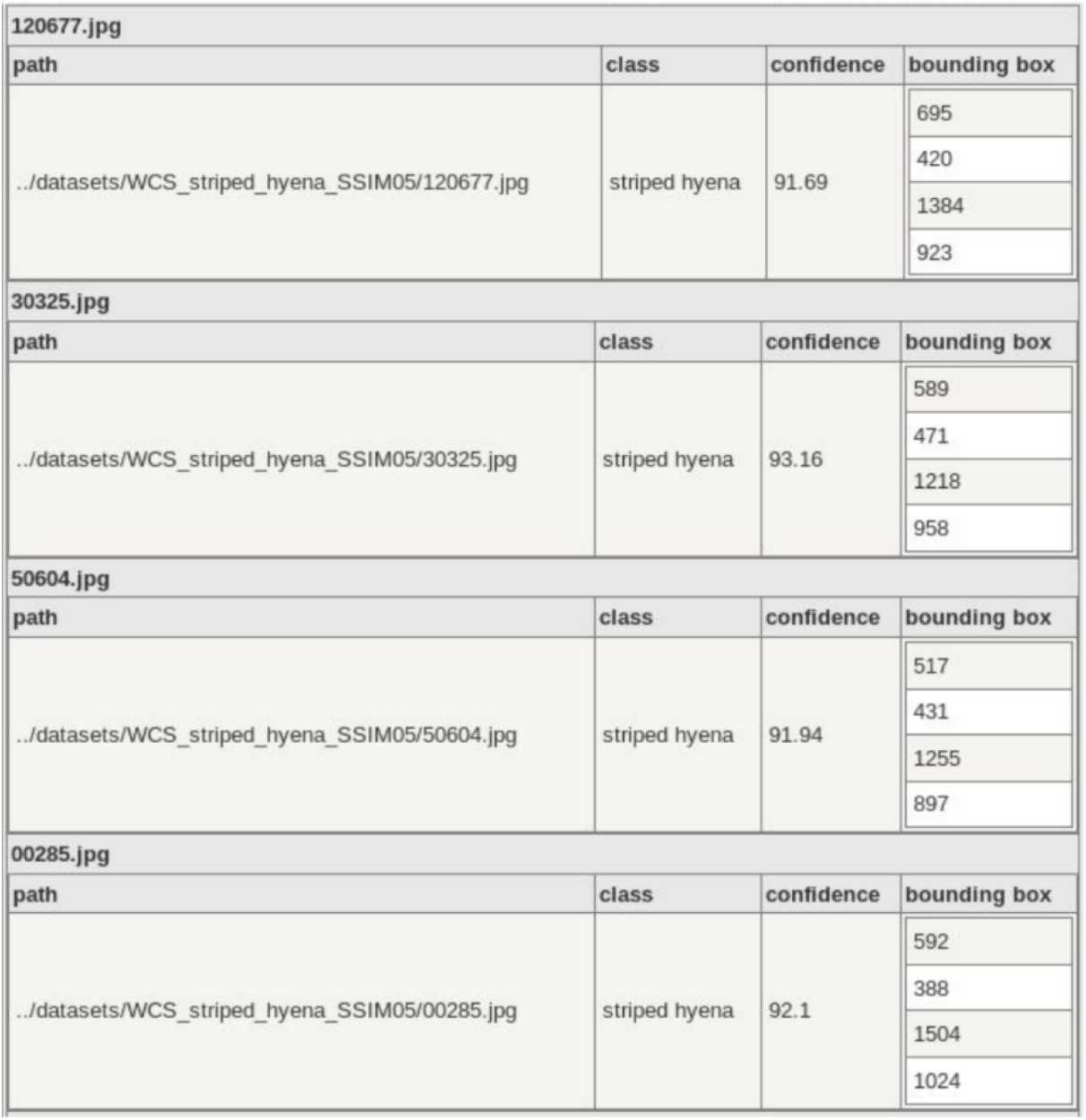
A sample of the Detailed Report, which shows per image data, including class label, classification confidence and bounding box data for each object. This preview is generated using json2table.com.

## V. Custom Model Training Using FlickR and Camera Trap Image Infusion

If the results attained by the default models are not sufficiently accurate for the user’s purposes, or the models are not sufficiently specific to the user’s domain, they may elect to custom train their own model or models. They may choose to use publicly available images such as FlickR and iNaturalist (FiN) images, infused with camera trap images as proposed by (Shepley, Falzon et al. 2020) or they may alternatively train using images from any source, including publicly available images, and/or their own trap images.

### a) Dataset and Classes

All datasets to be used for training should be placed in the *datasets* directory within the U-Infuse base directory. Users may add datasets of their choice to this directory or download U-Infuse FlickR annotated datasets from the U-Infuse GitHub page. The U-Infuse dataset repository contains 35 freely available single class datasets. Users may choose to use these datasets, or other publicly available datasets, or private datasets such as project specific camera trap images. User images are not shared, or accessible outside the user’s network or device, ensuring protection of intellectual property and privacy. All bounding box annotations must be placed in the *annotations* directory, also within the U-Infuse base directory. Users must ensure that any custom datasets are accompanied by corresponding annotation files, which must be placed in the *annotations* directory in a folder with the same name as the corresponding image dataset. For example, if the user adds the image dataset ‘*New England Jan*’ to *datasets* they must provide annotations in a folder named ‘*New England Jan*’ in the *annotations* directory. These annotations must be in PASCAL VOC format. Alternative annotation formats including YOLO may be supported in future releases. If a user does not have annotations, they may auto-annotate their images within U-Infuse, as discussed in Section 6.

Once the user has ensured that their image and annotation folders are located correctly in *datasets* and *annotations*, they may proceed to the training process, which can be initiated via the *Training Datasets and Classes* dialogue shown in Figure 5. Users must select one or more datasets for inclusion in training from the list of available datasets shown in the combo-box denoted by (1).

**Figure 5:**
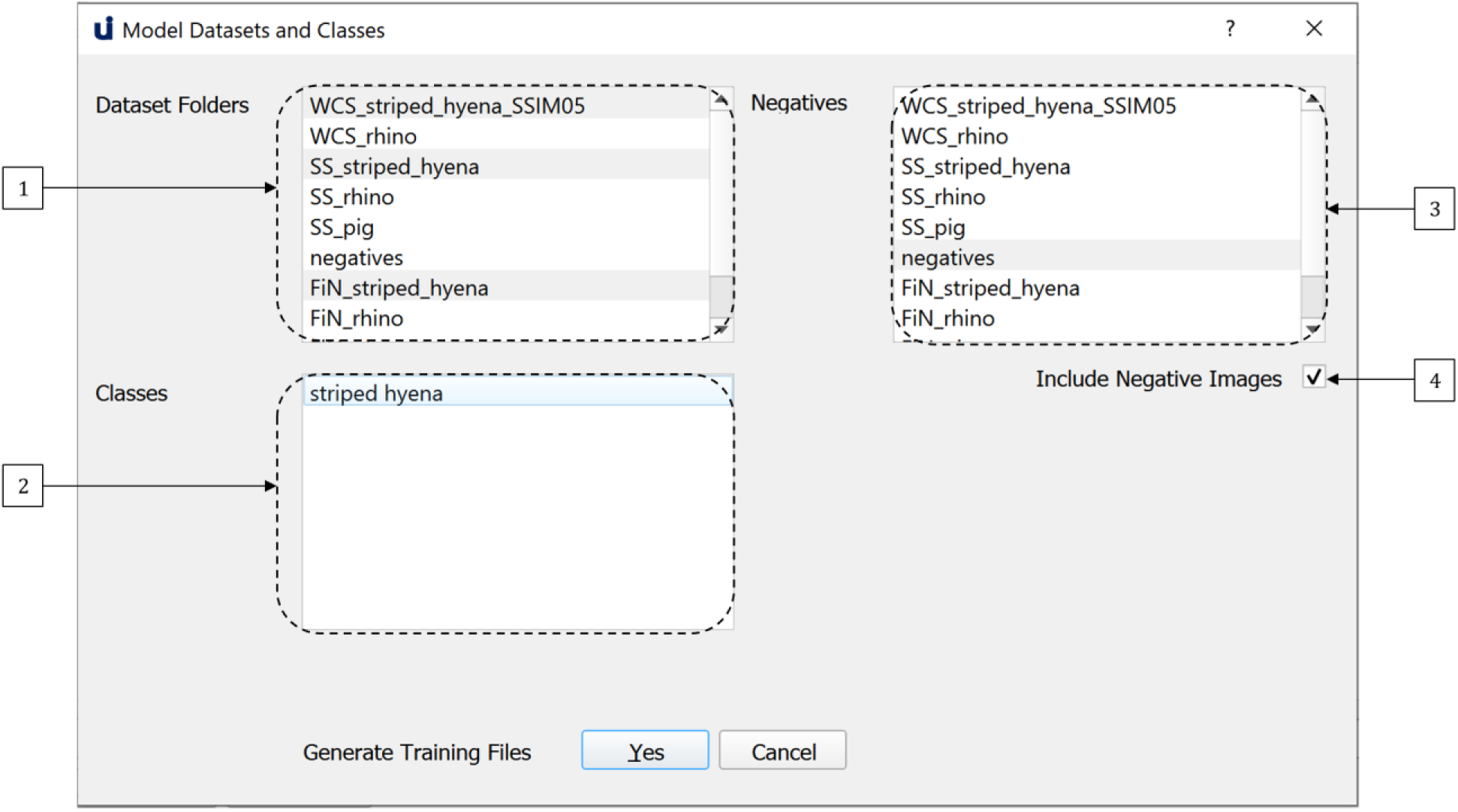
The Model Training dialogue allows users to custom train models on datasets and classes of their choice.

Once the user has selects one or more datasets, the list of classes available for training within those datasets is shown in the combo-box denoted by (2). If a dataset is deselected, classes specific to that dataset can no longer be chosen. U-Infuse supports both single class and multi-class training, meaning the user can select one or more classes for inclusion in training from (2). For example, the user may select five datasets, containing a total of twelve classes, but they may choose to only train on three of the available classes. They can then select one or more datasets from the combobox denoted by (3) for negative sampling. Alternatively, they can select the option to use all other classes for negative sampling. They may elect not to use negative sampling (4) however this is not recommended as negative sampling has been shown to significantly improve the ability of a model to discriminate between positive and negative objects (Tiwary 2018).

### b) Training

After dataset and class selection, the user may proceed to the training phase. Training requires the use of a Graphical Processing Unit (GPU) supporting CUDA (Nickolls, Buck et al. 2008) due to the computationally expensive nature of training deep neural networks. Deep learning is a resource-intensive process that cannot be effectively achieved using a standard CPU. GPUs are specialised processors used to process images, as they can handle large amounts of data, and support parallel processing. It is worth noting that U-Infuse does not currently support training without pre-initialised network weights, which requires thousands to millions of images to achieve acceptable results, due to random initialisation of weights. Instead U-Infuse provides capability for modifying pre-trained network weights using transfer learning, which requires comparatively minimal data and computational cost to develop effective deep learning models (Schneider, Taylor et al. 2018, Willi, Pitman et al. 2018).

U-Infuse features default training parameters established within our research program. They can be modified via the Model Training Settings Dialogue shown in Figure 6. Firstly, the user must select a pretrained model (1). U-Infuse provides 6 pretrained models, one of which can be selected as the backbone for further model training. User provided or trained models can also be used as the basis for further model training. Users may elect to modify the ‘Epochs’ (2) option. By default, training will proceed for 30 epochs, however users can increase or decrease the number of epochs, depending on the accuracy they seek. Reducing the number of epochs will reduce training time, however it may also reduce accuracy. Increasing the number of epochs means that the model is trained for longer, resulting in higher model training accuracy, but can lead to overfitting. Overfitting occurs when the network memorises the features of the training data, limiting its ability to generalise to other datasets. To avoid overfitting and allow monitoring of training, U-Infuse outputs data such as the training and validation loss, as well as Mean Average Precision (mAP) results calculated on a validation dataset after each training epoch.

**Figure 6:**
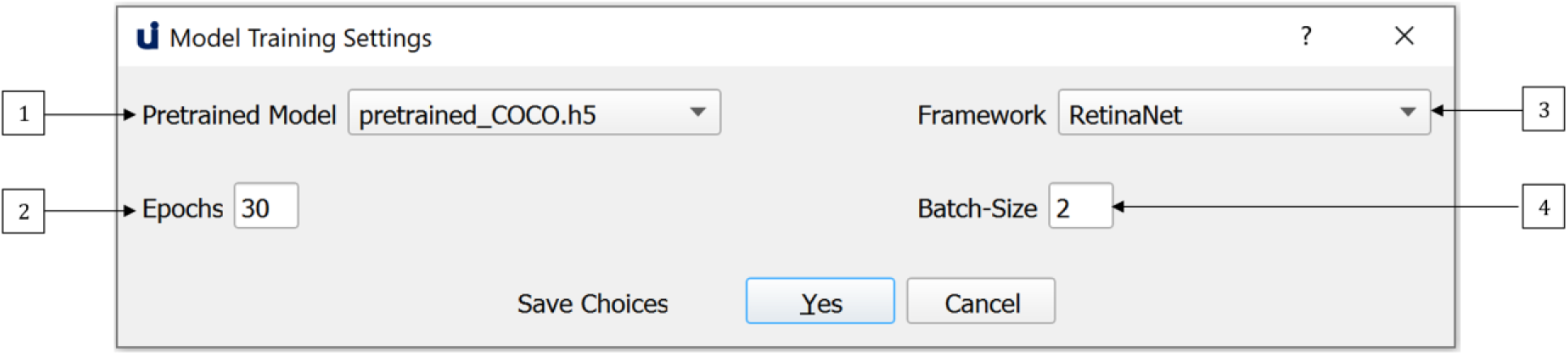
Default training parameters are provided however users may elect to modify these depending on their GPU capabilities and dataset size.

To use U-Infuse for training, users must have access to a GPU. As GPU capability may vary between users, the batch size used for training can be varied (4). A default batch size of 2 is provided, however users with high capability GPUs may elect to increase this value, while those with limited GPU resources may reduce the batch size to 1. The greater the batch size the more GPU memory required for training. Once the datasets, classes and training parameters have been selected, the user can proceed to generate the training scripts. Alternatively, the dialogue may be closed, with no changes saved by selecting the *Cancel* option. Any error or success messages are shown in the main frame output window.

If the training configuration process is successfully completed, the *Start Training* option on the main frame may be selected to train a RetinaNet model. Training will usually take several hours, depending on the dataset size, number of epochs and batch size used. During training the user will be updated by messages in the main frame, as shown in Figure 7. After each epoch, a snapshot of the model is saved in the *./snapshots* directory. A snapshot is a file containing the weights of the model after a given epoch and should not be confused with terms such as Snapshot Serengeti.

**Figure 7:**
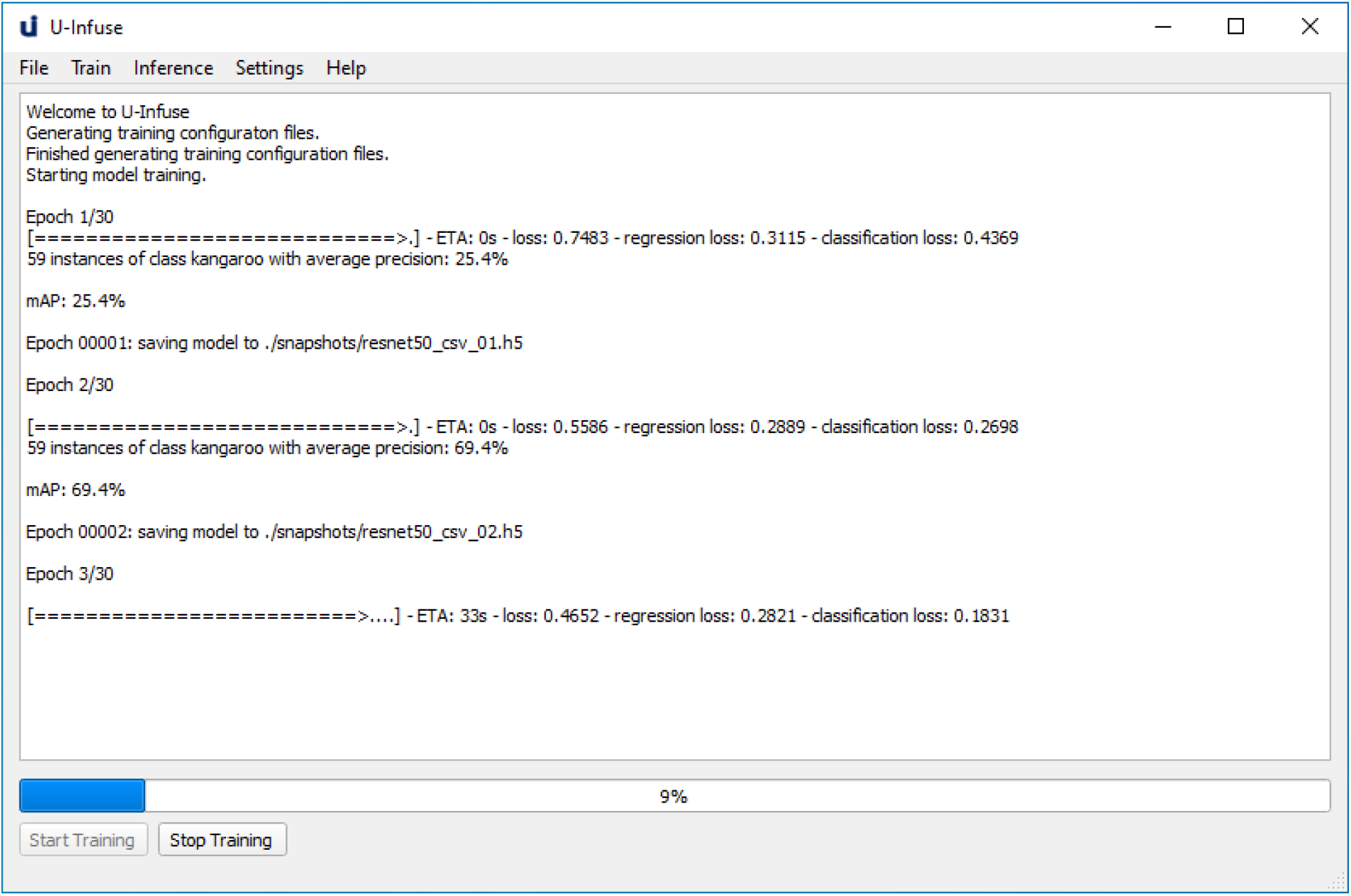
The main frame which allows users to start, stop and monitor training of custom models. Training progress can be monitored via the overall training loss, regression loss and classification loss. Generally, the lower the loss, the better the training. Model performance can also be monitored via the validation mAP, which is calculated on a subsample of the training data.

Once training is complete, the user can choose which snapshot they wish to retain as their final custom model. They can preview the performance of snapshots via the *Preview Custom Model* dialogue shown in Figure 8. They can choose snapshots via the dropdown list at (1), select their test images (2) and the number of images they wish to preview (3) as well as a level of accuracy (4). Once they run the model (5), they may name their model (6) and it (8). It is recommended that users delete all other models (7) as snapshots are large files. The exported model will be saved, to be reused via the *Object Detection* dialogue for object detection on any dataset, as described in Section 4.

**Figure 8:**
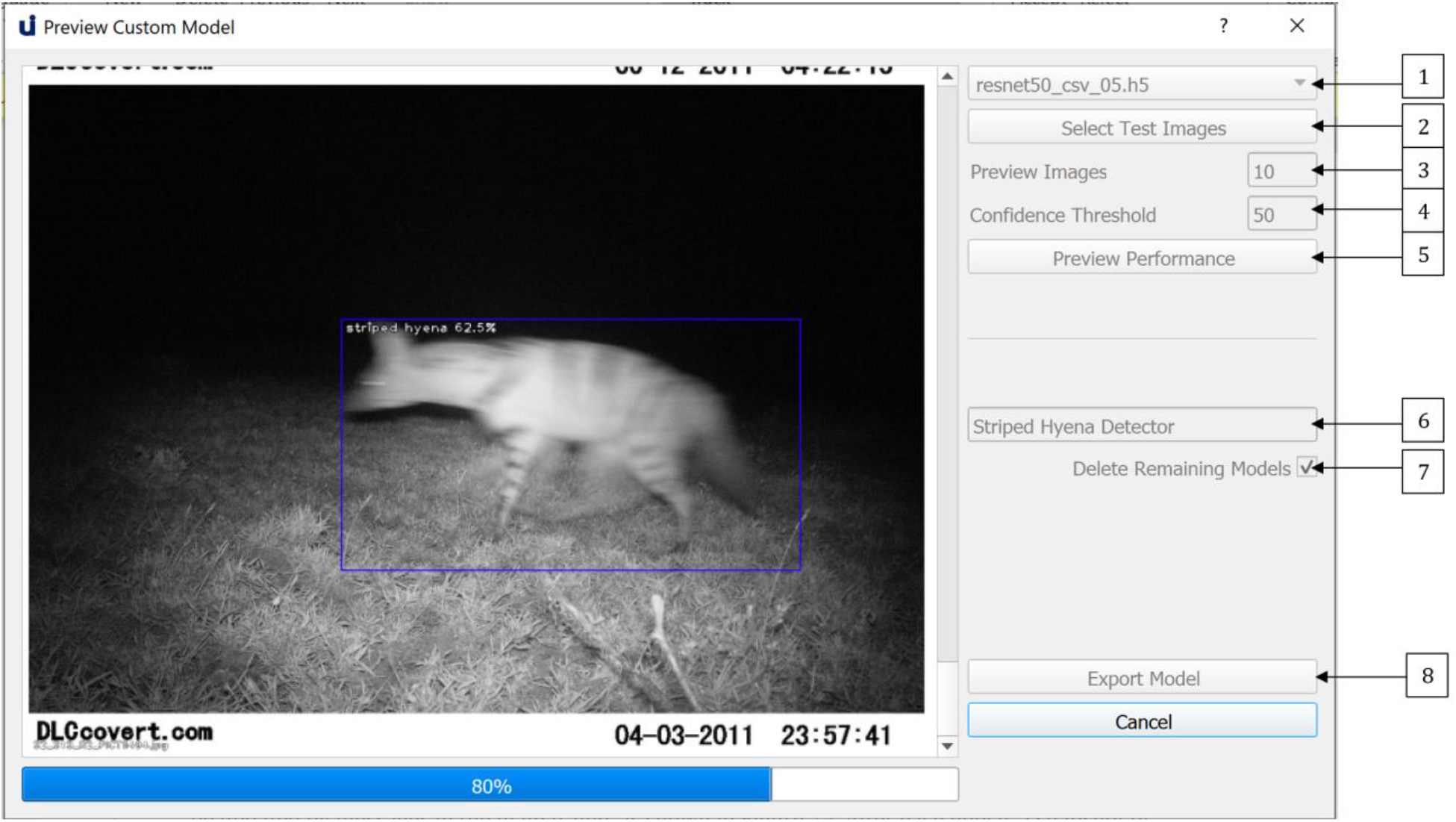
The Preview Custom Model dialogue allows the user to preview the performance of custom trained models on a small subset of randomly sampled test images. It also allows users to export chosen models for future use.

## VI Auto-Annotation and Manual Annotation Editing

One of the most time-consuming aspects of developing deep learning-based object detectors is the annotation of training images. U-Infuse automates this process, by allowing users to employ pretrained models, or a provided Single Class Annotator to annotate any number of images. This can be achieved via the *Auto-Annotation* dialogue shown in Figure 9. Users must add the folder/s containing the images they wish to annotate to the *datasets* directory. They can then access their chosen dataset via the dropdown list (1). If the user chooses to conduct multi-class annotation (2), they must select a pretrained model from the list of available options (4). The labels used to annotate objects will be chosen by the pretrained model. Alternatively, the user can provide a single class label (3) which will be used to annotate all bounding boxes generated by the provided auto-annotator. This is very useful in cases where the user wants to annotate a dataset containing objects for which they do not have an object detector. Users can also vary the confidence threshold (5). A higher confidence threshold means less bounding boxes are retained, while a lower confidence threshold allows more bounding boxes to be shown. Users may elect to show images (6) or not. Note, electing to show images is more computationally expensive, meaning the annotation process will usually take longer.

**Figure 9:**
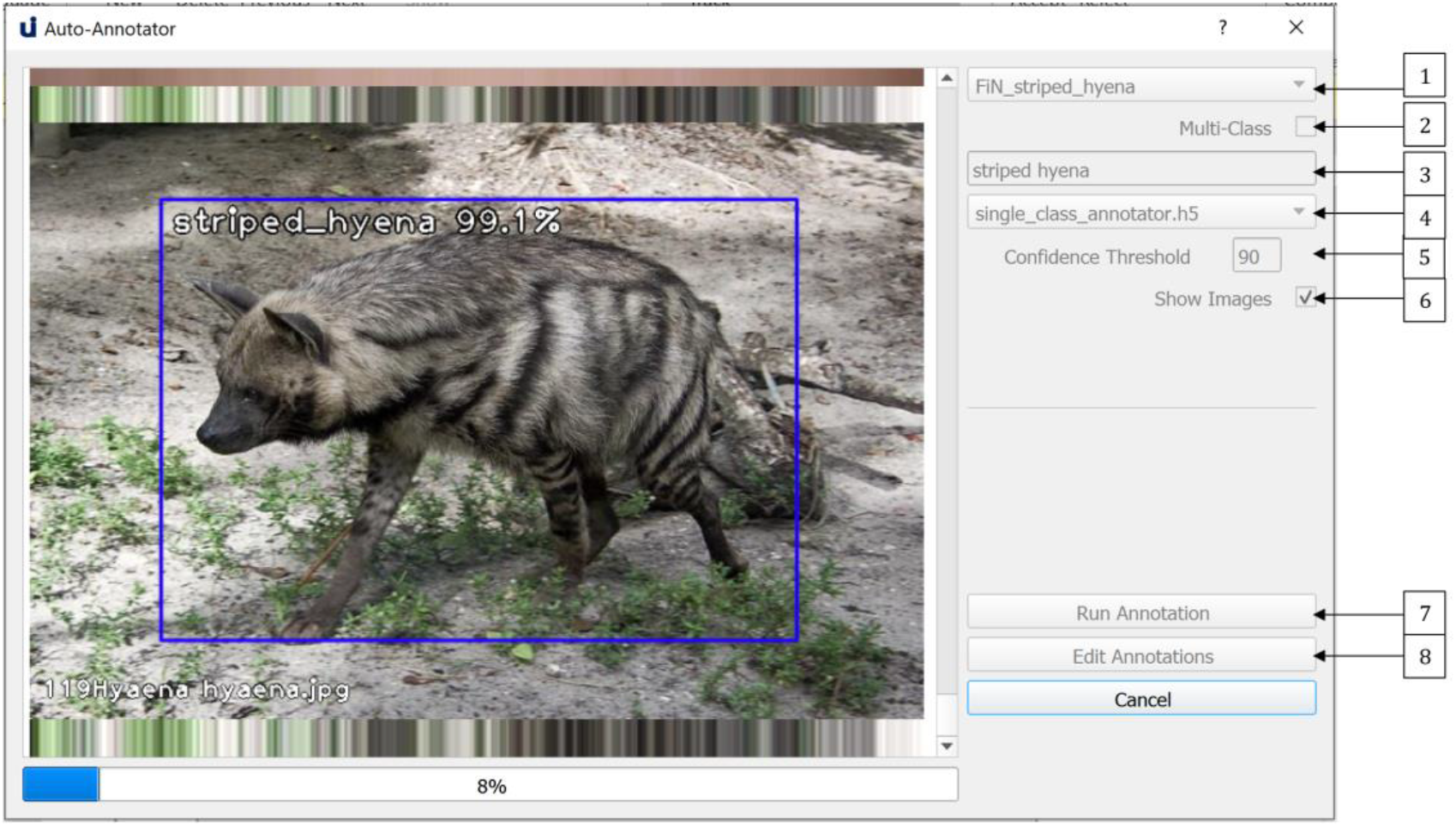
The Auto-Annotation dialogue allows users to achieve both single class and multi-class auto-annotation of images. Annotations can be edited via labelImg.

Once the user has run the annotation model (7), they may edit these annotations (8) via labelImg (Tzutalin 2015), to remove unwanted boxes, edit the position or size of boxes or add boxes around objects that were missed. Sometimes, the auto-annotator will miss objects, add extra boxes, or the boxes may not be optimal, for example, the trap image shown in Figure 10 contains an extra box which should be removed.

**Figure 10:**
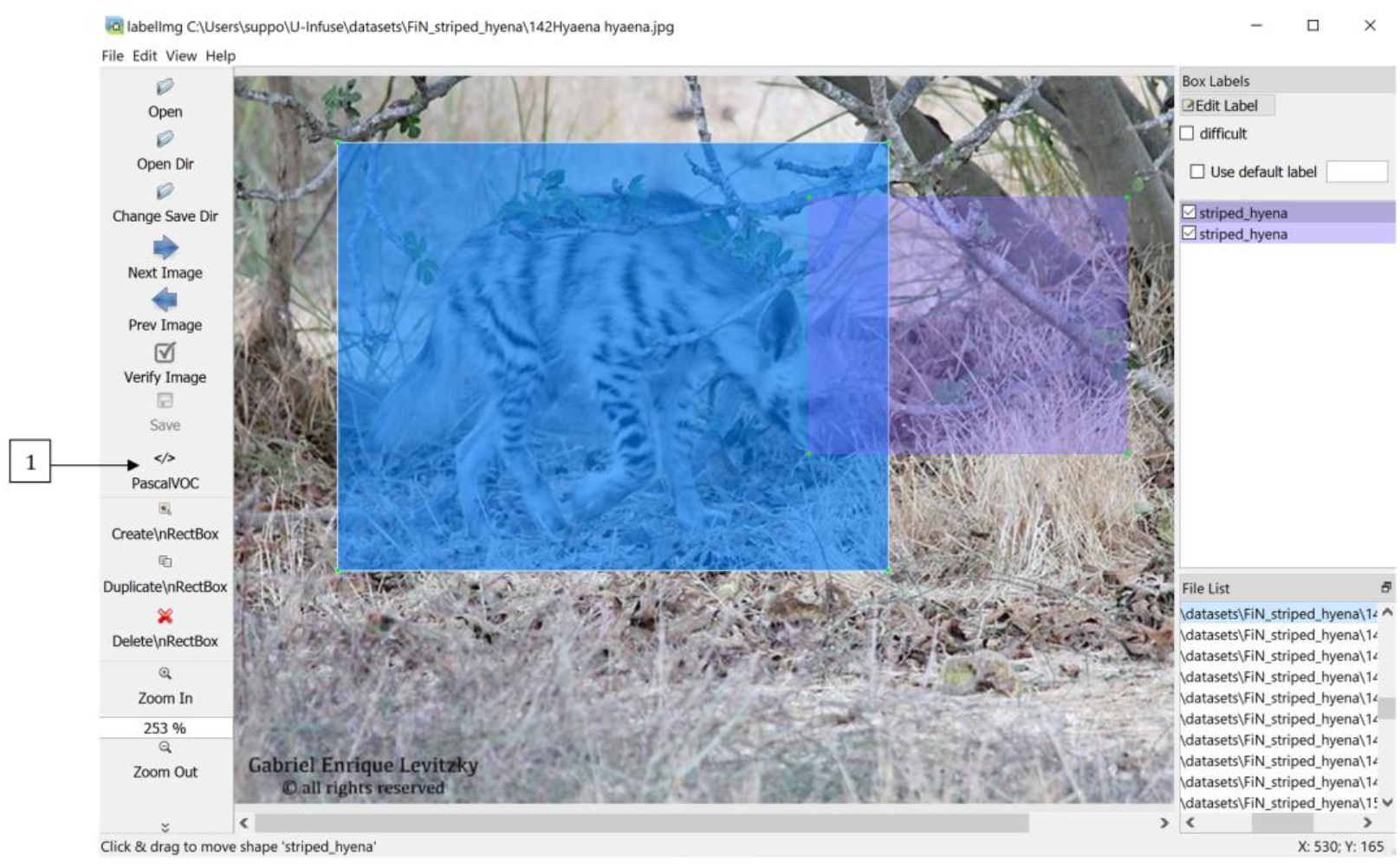
Annotations can be edited in labelImg. Bounding boxes can be added, removed or altered. Labels can also be altered.

Bounding boxes can be added, removed or resized, and labels can be changed within the labelImg GUI as shown in Figure 8. Users must ensure they save any altered annotations in Pascal VOC format by selecting the option denoted by (1) in Figure 10. Once this process is completed, the images and corresponding annotation files can be used to train models, as described in Section 5.

### Future Work and Conclusion

U-Infuse is an open source application which implements the location invariance methodology proposed by (Shepley, Falzon et al. 2020). It democratises deep learning and AI technologies by making deep learning more accessible for ecological practitioners. Furthermore, it can be used for location invariant object detection in fields outside of ecology. Its open source nature means members of the community can contribute to its development, incorporating it and using it to complement existing software and cloud-based platforms. Ecologists are encouraged to contribute their annotated camera trap images to the U-Infuse repository to contribute to the development of more powerful, location invariant object detectors.

A major constraint on the widespread deployment of AI in ecology and automated processing of camera trap images is the use of complex deep learning algorithms and processes, usually accessible only to computer scientists. U-Infuse bridges this gap by allowing ecologists to train their own models via a user-friendly GUI. U-Infuse is therefore an important connecting inter-phase between computer science and ecology as it allows field practitioners to undertake tasks usually only understood by and reserved to computer scientists. It represents a significant progression from early studies, which involved ecological practitioners collecting images, manually cataloguing and placing bounding boxes around animals, providing these images to computer scientists to develop domain specific object detectors, which is a major limitation of solutions such as ClassifyMe (Falzon, Lawson et al. 2020).

Furthermore, U-Infuse may be extended to support other frameworks including YOLOv3 (Redmon and Farhadi 2016), Faster RCNN (Ren, He et al. 2015) and Single Shot Detectors (Zhang, Wen et al. 2018). Incorporating FlickR API support to allow image downloading and sorting within U-Infuse, and integration of the SSIM image similarity tool and duplicate remover (Shepley, Falzon et al. 2020) would also extend its usability in the development of high performance, domain specific deep learning object detectors. Copying, distribution and modification of U-Infuse source code is encouraged. Accordingly, U-Infuse is distributed under the terms of a GNU General Public License (link).

## Data Accessibility

U-Infuse can be accessed and installed by visiting https://github.com/u-infuse/u-infuse. All datasets, and Jupyter notebooks/tutorials can also be accessed on this GitHub page.

## Notes

https://github.com/u-infuse/u-infuse/

### Competing Interest Statement

The authors have declared no competing interest.

