## Appendix for "U-Infuse: Democratization of Customizable AI for Object Detection"

### Appendix 1

| Model name | Classes |
|---|---|
| COCO_pretrained | 0: 'person', 1: 'bicycle', 2: 'car', 3: 'motorcycle', 4: 'airplane', 5: 'bus', 6: 'train', 7: 'truck', 8: 'boat', 9: 'traffic light', 10: 'fire hydrant', 11: 'stop sign', 12: 'parking meter', 13: 'bench', 14: 'bird', 15: 'cat', 16: 'dog', 17: 'horse', 18: 'sheep', 19: 'cow', 20: 'elephant', 21: 'bear', 22: 'zebra', 23: 'giraffe', 24: 'backpack', 25: 'umbrella', 26: 'handbag', 27: 'tie', 28: 'suitcase', 29: 'frisbee', 30: 'skis', 31: 'snowboard', 32: 'sports ball', 33: 'kite', 34: 'baseball bat', 35: 'baseball glove', 36: 'skateboard', 37: 'surfboard', 38: 'tennis racket', 39: 'bottle', 40: 'wine glass', 41: 'cup', 42: 'fork', 43: 'knife', 44: 'spoon', 45: 'bowl', 46: 'banana', 47: 'apple', 48: 'sandwich', 49: 'orange', 50: 'broccoli', 51: 'carrot', 52: 'hot dog', 53: 'pizza', 54: 'donut', 55: 'cake', 56: 'chair', 57: 'couch', 58: 'potted plant', 59: 'bed', 60: 'dining table', 61: 'toilet', 62: 'tv', 63: 'laptop', 64: 'mouse', 65: 'remote', 66: 'keyboard', 67: 'cell phone', 68: 'microwave', 69: 'oven', 70: 'toaster', 71: 'sink', 72: 'refrigerator', 73: 'book', 74: 'clock', 75: 'vase', 76: 'scissors', 77: 'teddy bear', 78: 'hair drier', 79: 'toothbrush' |
| Australian_Multi-class | 0: ‘bandicoot’, 1: ‘brushturkey’, 2: ‘bus’, 3: ‘car’, 4: ‘cat’, 5: ‘cow’, 6: ‘deer’, 7:’dog’, 8: ‘eagle’, 9: ‘echidna’, 10: ‘finch’, 11: ‘fox’, 12: ‘goanna’, 13: ‘goat’, 14: ‘hare’, 15: ‘horse’, 16: ‘kangaroo’, 17: ‘koala’, 18: ‘lyrebird’, 19: ‘magpie’, 20: ‘motorbike’, 21: ‘pademelon’, 22: ‘parrot’, 23: ‘person’, 24: ‘pig’, 25: ‘possum’, 26: ‘quoll’, 27: ‘rabbit’, 28: ‘train’, 29: ‘truck’, 30: ‘wallaby’, 31: ‘wallaroo’, 32: ‘wombat’ |
| pig_single_class | 0: ‘pig’ |
| striped_hyena_single_class | 0: ‘striped hyena’ |
| rhino_single_class | 0: ‘rhinoceros’ |
